# Modeling habitat suitability for insect pollinators in New York City: Two decades of change

**DOI:** 10.1101/2025.07.13.664483

**Authors:** Shira Linsk, Kristin Winchell, Anna Thonis

## Abstract

As cities change and expand, it is increasingly important to understand how urbanization is altering habitat suitability for wildlife, particularly insect pollinators. Using species distribution models (SDMs), we assessed spatial and temporal changes in habitat suitability for diurnal pollinators in New York City over two decades (2000s and 2010s), focusing on three insect orders: Lepidoptera, Hymenoptera, and Coleoptera. Our models revealed an 18.5% net decrease in pollinator habitat suitability citywide, with marked variation across boroughs, including localized increases in Manhattan and the Bronx. Climatic variables, especially temperature, solar radiation, and vapor pressure deficit, emerged as the strongest predictors of habitat suitability, while land cover changes had more localized effects. Among land cover types, urban forest was the only class to show increased suitability, suggesting that greening initiatives may buffer some climate-driven declines. These findings highlight the combined influence of climate and land use on pollinator habitat in cities, emphasizing the need to consider both factors when assessing urban biodiversity change. Understanding these dynamics is critical for designing urban greening initiatives that effectively support pollinator communities and inform conservation efforts in increasingly developed landscapes.

## Introduction

Urbanization describes the transformation of natural landscapes into developed spaces characterized by high human population density, expanded infrastructure, and vast areas of impervious surface. Research suggests that by 2030, approximately 5 billion people will reside in urban areas (Seto et al., 2012), intensifying pressure to convert more land into urban development. In the United States alone, urban land area is expected to increase from 3.1% to 8.1%, representing a ∼390,400 km^2^ increase in the amount of developed land (Nowak & Walton, 2000). The ongoing and projected rapid growth of urban areas poses a major threat to global biodiversity (Boakes et al., 2023). Cities introduce a range of novel stressors that can disrupt local ecological communities, altering ecosystem patterns, processes, and functions (Casanelles-Abella et al., 2021). Specifically, urban landscapes contain large areas of impervious surface and infrastructure, and also experience elevated levels of noise, light, and air pollution—all of which can negatively affect local flora and fauna. Several studies have found that these types of chronic sensory pollutants disrupt urban wildlife by altering phenology, inducing physiological stress, shifting migratory behavior, and increasing mortality rates (Candolin, 2024; Kok et al., 2023).

Other studies have identified habitat availability as the primary driver of urban wildlife declines (Morpurgo et al., 2024; Jain et al., 2021). In sum, there is widespread agreement among ecologists that urbanization is a powerful driver of global biodiversity declines (Simkin et al., 2022; Piano et al., 2020; Tscharntke & Batáry, 2023; McKinney, 2004). However, a growing body of research suggests that urban areas can actually function as ecological hotspots harboring rare species (Boakes et al., 2024) and, when well maintained, supporting complex ecological communities.

As urban development transforms natural landscapes, urban green spaces are becoming increasingly critical for the preservation of native species (Goddard et al., 2010). Urban wildlife can benefit from urban infrastructure by exploiting anthropogenic resources commonly found in cities, such as garbage and other waste (Baker & Harris, 2007), or intentional sustainable infrastructure such as green roofs and community gardens. Furthermore, cities can serve as migratory stopovers for birds (Spotswood et al., 2021) and refuges for other native wildlife. For example, the common rain frog (*Breviceps adspersus*) in Britain has maintained stable urban populations even as their numbers decline in rural areas (Carriera & Beebee, 2002). Similarly, Hagen et al. (2017) found that of 529 bird species, 12.5% were found only in urban areas.

Furthermore, these urban-dwelling birds were primarily native to the study region (e.g., burrowing owls, black-and-rufous warbling finches, bronzed cowbirds), in contrast to more commonly recognized non-native urban species such as feral pigeons. With respect to arthropods, McFrederick and LeBuhn (2006) documented higher bumble bee abundance in the city of San Francisco than in neighboring suburbs. These findings all highlight the capacity of urban environments to support a diversity of species, underscoring the importance of understanding how habitat suitability for urban wildlife is changing over time.

As urban habitats continue to change, it is essential to consider the implications for ecosystem services that urban-dwelling organisms provide (Weiskopf et al., 2024). Insects, in particular, support key functions such as pollination, decomposition, and nutrient cycling (Vanbergen et al., 2013, Schowalter et al., 2018; Uma et al., 2023). A substantial proportion of insect species function as pollinators, spanning a wide range of taxonomic groups such as bees, butterflies, beetles, and flies (Requier et al., 2023; Rader et al., 2020). Approximately 87.5% of the world’s flowering plant species are pollinated by animals, primarily insects (Hoshiba & Sasaki, 2008), meaning the ongoing and rapid decline of insect populations is expected to drive the loss of countless plant species (Hallmann et al., 2017; Sánchez-Bayo and Wyckhuys, 2019). Insects are also integral to urban food webs as a vital food source for birds, amphibians, and mammals (Hallmann et al., 2017). Beyond their ecological roles, many insect species also have economic value through apiaries and biological control in agriculture (Rust & Su, 2012), with insect-mediated ecosystem services estimated at $57 billion annually in the United States alone (Losey & Vaughan, 2006). Widespread losses of pollinator diversity threaten a range of ecological and economic processes, with cascading effects on species interactions, plant pollination, and habitat integrity (Christmann et al., 2019). Therefore, preserving a wide diversity of pollinator species is critical for maintaining both agricultural productivity and ecological resilience (Ollerton, 2017). While crop pollination often receives the most attention, non-crop pollination services also play a key role in supporting biodiversity and overall ecosystem health, highlighting the importance of conservation efforts that extend beyond agricultural systems. As urbanization continues to accelerate, conserving insect pollinators must remain a central priority, and city-level monitoring is a critical first step toward effective conservation (Sanllorente et al., 2025).

To guide these efforts, it is essential to understand how urbanization shapes habitat suitability for pollinators. Despite the critical role of pollinators in urban ecosystems, relatively little is known about how their habitat suitability is changing in response to urbanization. Much of the existing research on urban biodiversity has focused on vertebrates or medically significant insects, with limited attention given to other communities across space and time (Casanelles-Abella et al., 2021; Titley et al., 2017; Mammola et al., 2021, Collins et al., 2024). In particular, few studies have assessed the habitat suitability of pollinators in cities or how those patterns vary across taxonomic groups. To address this gap, we examine how habitat suitability for diurnal insect pollinators in New York City has changed over the past two decades. Using species distribution models (SDMs), tools that combine species occurrence data with environmental variables to estimate habitat suitability (Elith & Leathwick, 2009), we assess spatial and temporal trends in pollinator habitat and identify the environmental characteristics associated with higher suitability. By evaluating shifts in habitat suitability across space, time, and insect taxa (i.e., Lepidoptera, Hymenoptera, Coleoptera), our study provides a valuable baseline for long-term monitoring of urban insect communities. We identify where, and in what direction, pollinator habitat suitability has changed in NYC and offer insight into the environmental factors potentially shaping urban insect distributions.

## Methods

### Study extent

Our study area included all five boroughs of New York (Manhattan, the Bronx, Queens, Brooklyn, Staten Island), a highly urbanized environment with the highest human population in the United States (8.2 million in 2023) spread across 709 km^2^. It falls within the temperate deciduous forest biome, characterized by warm summers with the highest temperatures and peak rainfall occurring during the summer months.

### Occurrence records

We downloaded all occurrence records for the insect orders Lepidoptera, Hymenoptera, and Coleoptera in New York City from 2000 to 2019 (inclusive) from the Global Biodiversity Information Facility (GBIF). After manually removing occurrences located in water, 19,136 observations remained in the dataset. To further refine the dataset, we excluded all records with coordinate uncertainty greater than 500 meters and used the clean_coordinates function from the CoordinateCleaner R package (Zizka et al., 2017) to identify and remove incorrectly georeferenced occurrences. We selected an uncertainty threshold of 500 m to ensure the spatial uncertainty of occurrence records did not exceed the resolution of the environmental raster layers (i.e., 1 km). Incorrectly georeferenced records included those with identical latitude and longitude, coordinates recorded as (0,0), and locations corresponding to biodiversity institutions such as museums. This reduced the dataset to 17,169 records. Finally, we excluded all records corresponding to species not known to engage in pollination. For Lepidoptera, we retained all butterfly families, including Hesperiidae, Riodinidae, Lycaenidae, Pieridae, Papilionidae, and Nymphalidae. All Hymenoptera families were initially retained, but we then excluded those unlikely to participate in pollination, such as gall wasps, ants, wood-boring sawflies, and non-nectar-feeding parasitoids. Specifically, we removed the following families: Ampulicidae, Chalcididae, Cynipidae, Diprionidae, Evaniidae, Formicidae, Pelecinidae, Rhopalosomatidae, Siricidae, and Tenthredinidae. Among Coleoptera, we retained only families associated with floral visitation or known indirect pollination (Soares et al., 2022; Mawdsley, 2003; Camurça et al., 2024; Traylor et al., 2023; Herrera, 2021; deMaynadier et al., 2024), specifically: Cantharidae, Cerambycidae, Chrysomelidae, Melyridae, Mordellidae, Nitidulidae, and Scarabaeidae (Figure 1). Diptera were excluded from the analysis due to limited GBIF data availability. This further reduced our dataset to 12,023 observations.

**Figure 1.**
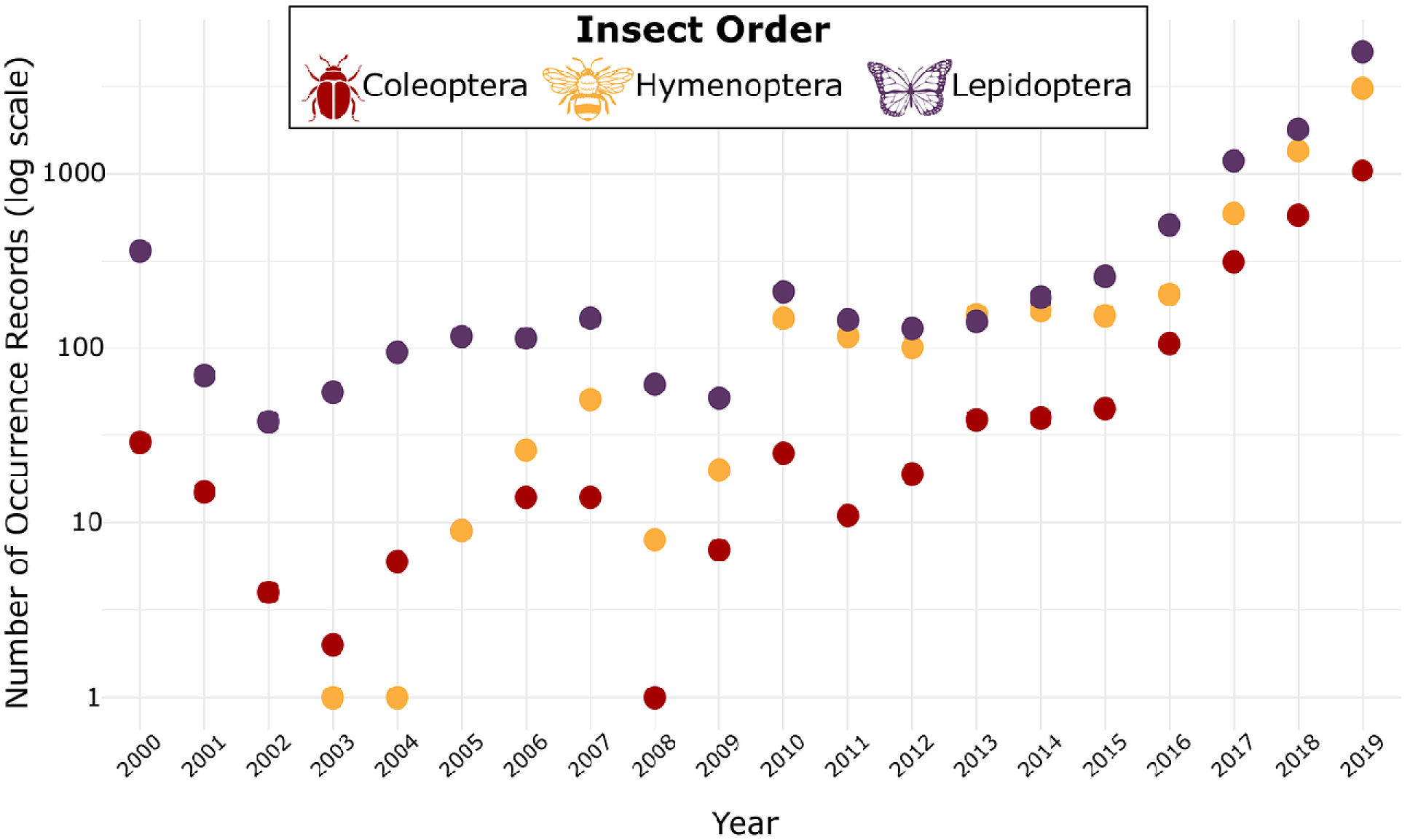
Occurrence records for the focal insect orders—Coleoptera, Hymenoptera, and Lepidoptera— obtained from GBIF for the period 2000–2019.

For decadal analyses, the 12,023 records were divided into two time periods: 2000–2009 (474 records) and 2010–2019 (11,548 records). After removing duplicate occurrences, we retained 156 unique records for the 2000s and 10,589 for the 2010s. Due to the limited sample size in the 2000s dataset, we did not apply spatial or environmental thinning (Fourcade et al. 2014). In contrast, the larger sample size and evident spatial bias in the 2010s dataset required additional processing. To reduce spatial sampling bias in the 2010s dataset, we applied kernel density estimation (KDE) to the occurrence records and thinned them by probabilistically subsampling points inversely proportional to local sampling density (Verbruggen et al. 2013, Fourcade et al. 2014, Botella et al. 2019, Baker et al. 2024). Specifically, we used a 2D KDE over a 1000×1000 grid and selected 5,000 presence records, assigning lower sampling probabilities to occurrences in densely sampled areas. This KDE-based approach can be more effective than spatial distance thinning because it accounts for continuous variation in sampling effort rather than applying often arbitrary distance thresholds, allowing for more nuanced bias correction in unevenly surveyed landscapes (Steen et al. 2021, Baker et al. 2024).

### Environmental predictor variables

We downloaded monthly climate data from CHELSA (Karger et al., 2017) at a resolution of 1 km (30 arcseconds) for 2000–2019 (inclusive). Variables included mean daily air temperature, mean daily minimum and maximum 2-meter air temperature, surface downwelling shortwave radiation, and vapor pressure deficit. For each variable, annual means were calculated and then averaged across each decade (i.e., 2000–2009, 2010–2019) for use in subsequent analyses. These specific climatic variables were selected to capture the primary environmental factors believed to be influencing pollinator habitat suitability (Elith & Leathwick, 2009). As ectotherms, insects rely on external conditions to regulate their body temperature, making them particularly sensitive to temperature changes. Temperature directly affects metabolic rates and behavior (Abdullah, 1961). Minimum daily temperatures are especially relevant for understanding overwintering success, diapause timing, and crepuscular activity (Roe et al., 2024; Chen & Seybold, 2014). In contrast, maximum daily temperatures constrain foraging behavior, as many pollinators have upper thermal limits beyond which activity and performance decline (Johnson et al., 2023).

Temperature also influences floral phenology and bloom periods, which in turn determine the availability of nectar and pollen resources (Scaven & Rafferty, 2013). Surface downwelling shortwave radiation serves as a proxy for sunlight availability, with higher values generally associated with increased pollinator activity and visitation to urban green spaces (Watson et al., 2022). We also included vapor pressure deficit, a measure of atmospheric dryness, as it can increase desiccation risk and contribute to vegetative stress, both of which may affect pollinator abundance and behavior.

In addition to climatic variables, we incorporated spatially explicit data on land cover, human population density, and air pollution. Land cover data were downloaded from the National Land Cover Database (NLCD; USGS, 2024) on decennial years at a spatial resolution of 30 m. Although the NLCD defines 16 distinct land cover classes, we reclassified these into six broader land cover classes most relevant to insect ecology. Specifically, The final six land cover classes were: (1) impervious surface cover, including developed land (low to high intensity) and barren areas; (2) forest cover, comprising evergreen, deciduous, and mixed forests; (3) herbaceous cover, including grasslands, sedge, shrubland, and open space; (4) cultivated land, such as croplands and pastures; (5) wetlands, both woody and herbaceous; and (6) open water.

We downloaded human population data from the National Historical Geographic Information System (NHGIS, Manson et al., 2024) for the years 2000, 2010, and 2020. Then we spatially joined population data with an NYC shapefile containing city boundaries to merge by borough. We calculated population density in R using the land area (km²) of each borough then averaged the population density for each decennial year and rasterized the results into two decadal files.

In addition, we included fine particulate matter (PM2.5), an air pollutant composed of particles less than 2.5 micrometers in diameter, as an air quality variable in our models due to its known ecological and health impacts. We retireived daily PM2.5 data via the Environmental Protection Agency’s API for NYC’s five boroughs (Manhattan, Brooklyn, Queens, Bronx, and Staten Island) for the years 2000 to 2020 (EPA, 2025). Using the he daily air quality data (i.e., PM2.5) we quantified the decadal averages per borough and then rasterized to represent the averages per borough over each decade.

We processed all environmental predictor variables to match in spatial resolution (1 km), geographic extent (New York City boroughs), and coordinate reference system (WGS84). We aggregated environmental predictor variables by decadal period (2000–2009, 2010–2019) to ensure a consistent temporal extent across datasets with varying temporal resolutions (e.g., annual vs. monthly). We resampled continuous variables using bilinear interpolation, while we used the nearest-neighbor method to resample categorical variables (e.g., NLCD land cover).

### Background point generation

To reduce sampling bias and improve model discrimination we generated background points using a target-group background (TGB) approach combined with environmental stratification. The TGB method selects background points from occurrence records of taxa assumed to share similar sampling biases with the focal taxa. In our case, background points were drawn from insect orders other than Lepidoptera, Hymenoptera, and Coleoptera—our three focal groups—under the assumption that non-focal insect taxa were collected under comparable survey conditions (Phillips et al., 2009; Barber et al., 2021). To construct the TGB pool, we first generated a large set of candidate background points using a kernel density bias layer created from all insect occurrence records within each decade. This ensured that background sampling reflected the spatial structure of collection effort. We then excluded all occurrences associated with the focal orders, leaving only non-focal insect taxa in the TGB pool.

To ensure equal prevalence between presence and background data, we generated the same number of background points as presence points for each model, following the recommendations of Barbet-Massin et al. (2012). To further improve model performance, we applied environmental stratification to the TGB points. For each background candidate point, we extracted environmental values and computed its multivariate Euclidean distance to the centroid of the environmental conditions at the species’ presence locations. We then excluded the 10% of background points most similar to the presence data in environmental space (Varela et al., 2014), thereby reducing overlap between presence and background environments. This approach enhances the model’s ability to distinguish suitable from unsuitable conditions by increasing contrast between presence and background points and mitigating the influence of shared environmental biases. Occurrence data were binned by decade, and both the TGB pool and environmental stratification were restricted to records from the same decade, ensuring temporal consistency in background sampling.For the 2010s order-specific models, background points were generated separately for Lepidoptera, Hymenoptera, and Coleoptera. In each case, the TGB pool excluded only the focal order, retaining occurrence records from the remaining insect taxa in New York City. This order-specific TGB approach ensured that background points reflected the sampling bias of related but non-focal groups while maintaining temporal consistency by limiting all background generation to 2010–2019 occurrence data.

### Random Forest models

To evaluate species-environment relationships, we implemented a Random Forest (RF) classification approach using 100 replicate model runs. For each iteration, occurrence data were randomly split into training (70%) and testing (30%) subsets. RF models were trained using 500 trees (ntree = 500), with the number of variables considered at each split (mtry) set to the square root of the number of predictor variables plus one, and a minimum terminal node size of five (Probst et al. 2018). Model performance was assessed on the withheld test data using the area under the receiver operating characteristic curve (AUC), sensitivity (true positive rate), specificity (true negative rate), the true skill statistic (TSS = sensitivity + specificity – 1), and Cohen’s Kappa. Thresholds for binary classification were selected using Youden’s J statistic, which maximizes the sum of sensitivity and specificity (Liu et al. 2005, Jiménez-Valverde and Lobo, 2007). Performance metrics were only computed for iterations in which both presence and background classes were represented in the test set. This iterative framework allowed for robust evaluation of model stability and performance across repeated subsampling. These methods were applied to both the aggregated models incorporating all focal orders (Lepidoptera, Hymenoptera, and Coleoptera) and the 2010s order-specific models.

To assess variable importance, we extracted the MeanDecreaseAccuracy metric from each RF model and summarized it across runs. This metric reflects the decrease in model accuracy when a given predictor is randomly permuted, providing an estimate of that variable’s contribution to predictive performance (Breiman 2001, Cutler et al. 2007). We also generated partial dependence plots (PDPs) to visualize the marginal effect of each predictor on predicted habitat suitability, while holding all other variables constant. PDPs essentially show how changes in a single variable influence model predictions, helping to interpret the direction and shape of species-environment relationships (Elith et al. 2008).

## Results

### Model performance

We generated five model types: two decade-specific models that included all focal insect orders (2000s and 2010s), and three models from the 2010s focused individually on Hymenoptera, Lepidoptera, and Coleoptera. All models demonstrated strong predictive performance (Table 1). Both the 2000s and 2010s aggregated models performed well overall, but the 2000s models exhibited greater variability across iterative runs, particularly in AUC scores (SD = 0.03 for 2000s vs. 0.003 for 2010s). Variation in other performance metrics, however, was comparatively minor between decades.

**Table 1.** Model performance metrics for decade-specific models combining all three focal orders (Hymenoptera, Lepidoptera, and Coleoptera) and the order-specific models for the 2010s. Metrics include Area Under the Curve (AUC), True Skill Statistic (TSS), Sensitivity, Specificity, and Cohen’s Kappa.

| Model | AUC | TSS | Sensitivity | Specificity | Kappa |
| --- | --- | --- | --- | --- | --- |
| All orders – 2000s | 0.89 | 0.72 | 0.78 | 0.93 | 0.78 |
| All orders – 2010s | 0.95 | 0.78 | 0.87 | 0.91 | 0.77 |
| Hymenoptera – 2010s | 0.94 | 0.77 | 0.87 | 0.88 | 0.77 |
| Lepidoptera – 2010s | 0.93 | 0.72 | 0.84 | 0.88 | 0.72 |
| Coleoptera – 2010s | 0.88 | 0.60 | 0.77 | 0.83 | 0.60 |

All order-specific models for the 2010s exhibited strong predictive performance (Table 1). Models for Hymenoptera and Lepidoptera performed similarly well across all metrics, with particularly high AUC, sensitivity, and specificity values. While still performing adequately, the Coleoptera model showed comparatively lower values across metrics, especially for TSS and Kappa, indicating reduced discriminatory power relative to the other two orders. Variation across iterative model runs was relatively low for all 2010s order-specific models, though some differences were evident between taxa (Table 1). The Coleoptera model exhibited the highest variability across all performance metrics, particularly for specificity, sensitivity, and TSS. In contrast, the Hymenoptera and Lepidoptera models showed more consistent performance, with markedly lower standard deviations across metrics, especially for AUC and Kappa.

### Habitat suitability

The aggregated models (2000s and 2010s) revealed an overall decline in habitat suitability across the study area over time. Mean suitability decreased from 0.27 (SD = 0.17, Figure 2a) in the 2000s to 0.22 (SD = 0.24, Figure 2b) in the 2010s, representing an overall decrease of 18.5% suitable habitat. Spatial trends in suitability shifts were highly variable across the five boroughs of New York City. While the overall trend showed a decline in habitat suitability between the 2000s and 2010s, several boroughs exhibited localized increases. Manhattan showed the largest increase in suitability with a mean difference of +0.234 (71.4%) and the Bronx showed a marginal increase of +0.020 (9.5%). In contrast, Brooklyn and Queens experienced substantial declines in habitat suitability, with a mean difference of –0.113 (30.8%) and –0.108 (46.8%), respectively. Staten Island also declined, though more modestly, with a mean decrease of –0.026 or 10.4% (Figure 2c). Mean suitability in the 2010s did not differ substantially across the three focal insect orders, based on results from the order-specific models (Figure 3). Spatial trends in suitability are closely aligned with patterns observed in the combined aggregated model for the same period.

**Figure 2.**
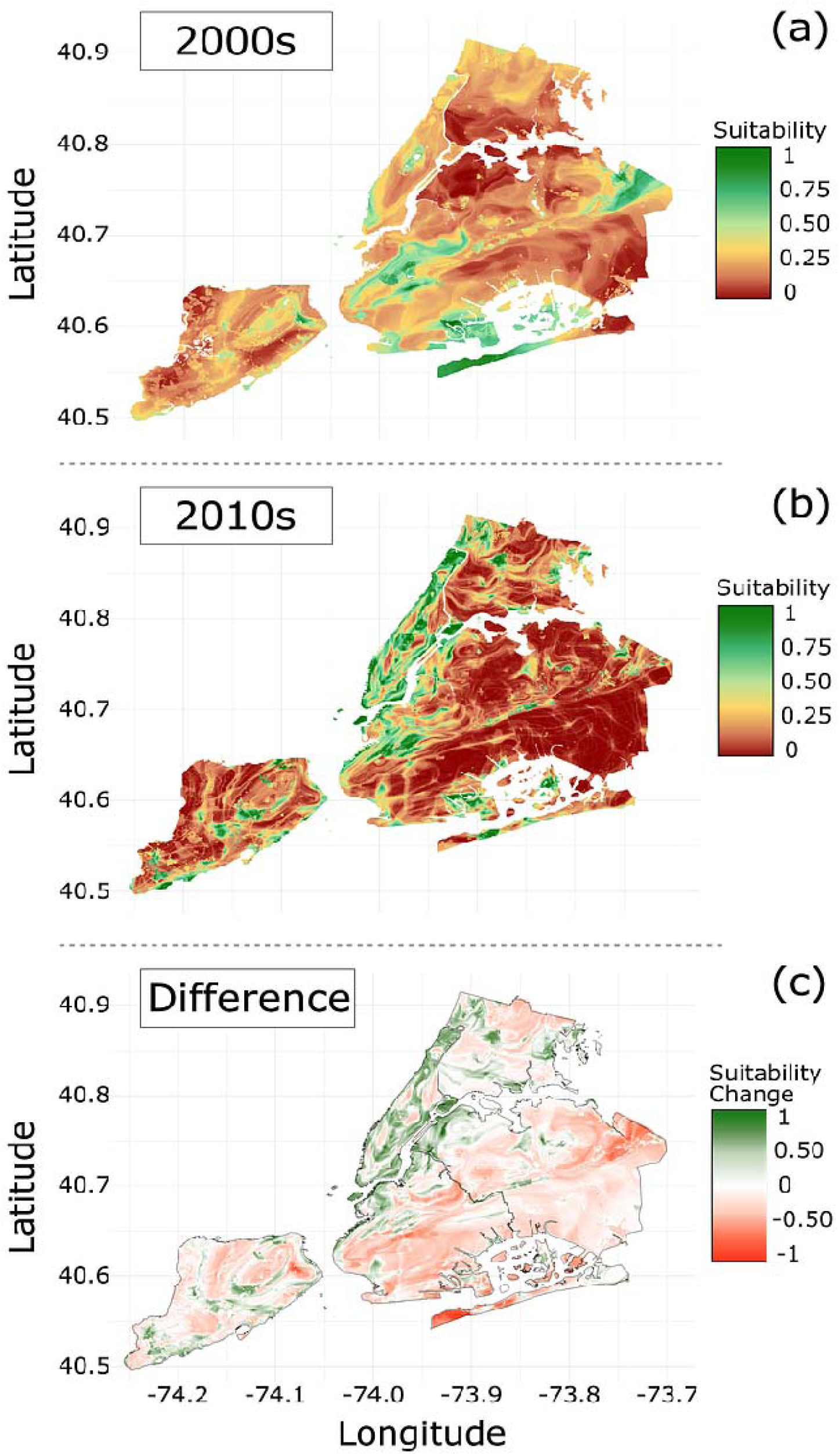
(a) Predicted habitat suitability for diurnal pollinators in New York City for the 2000s; (b) for the 2010s; and (c) the difference in mean habitat suitability between decades, with positive values (green) indicating areas that became more suitable over time and negative values (red) indicating areas of decline.

**Figure 3.**
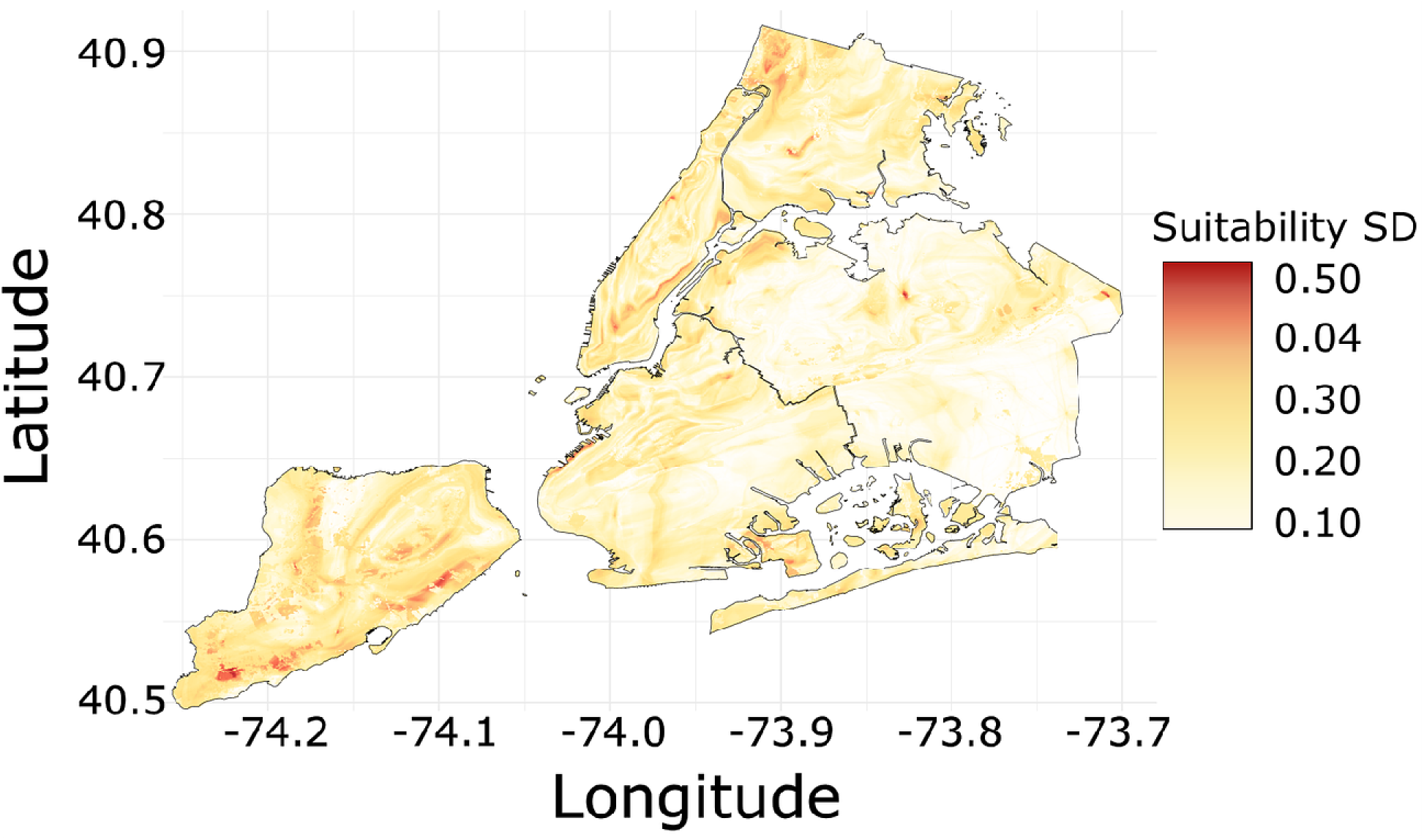
Standard deviation in modeled habitat suitability across three focal orders for the 2010s.

### Variable importance and direction of effect

Across both decades, the aggregated models for the 2000s and 2010s identified minimum temperature, maximum temperature, vapor pressure deficit, mean temperature, and surface downwelling shortwave flux as the top five most important predictors of habitat suitability, based on the mean decrease in accuracy (Figure 4). While the same set of variables ranked highest in the order-specific models, their relative importance shifted between decades. For instance, vapor pressure deficit was the top predictor in the 2010s, but ranked fourth in the 2000s, whereas minimum temperature consistently remained among the top three. In contrast, air pollution, human population density, and land cover consistently showed low predictive power in both the 2000s and 2010s models. Interestingly, mean air pollution was more important than maximum air pollution across models. Evaluation of habitat suitability by land cover classification revealed notable changes between decades: impervious land cover (median difference = –0.118) and wetlands (median difference = –0.141) experienced the largest decreases in suitability, while urban forest was the only class to show an increase (median difference = 0.147) (Figure 5).

**Figure 4.**
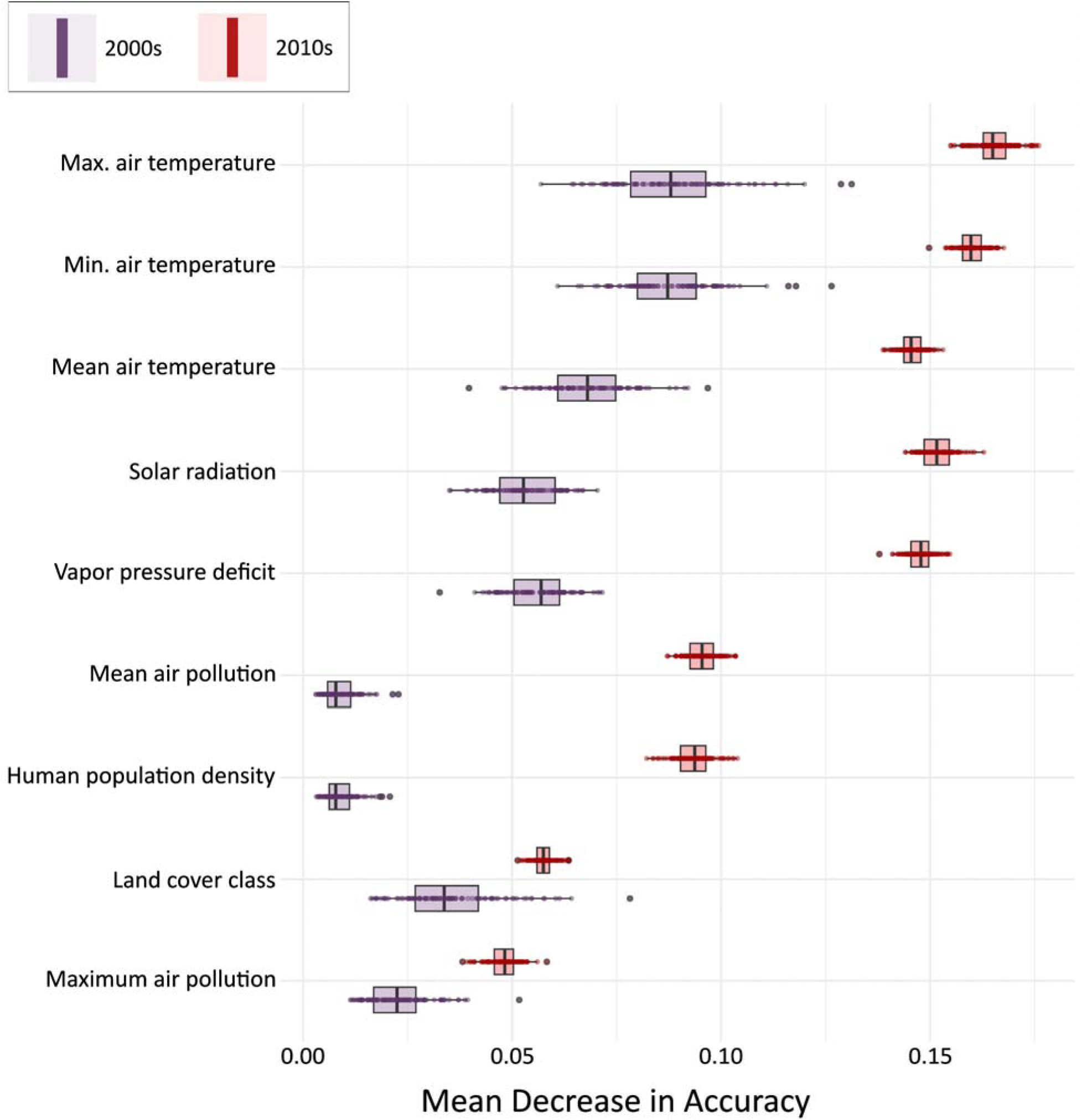
Box-and-whisker plots of 2000s (purple) and 2010s (red) variable importance. Variables are ordered by mean decrease in accuracy across both models. Each box represents the interquartile range (IQR), with the median indicated by a horizontal line. Whiskers extend to values within 1.5 times the IQR, highlighting the variability in predictor importance across model runs.

**Figure 5.**
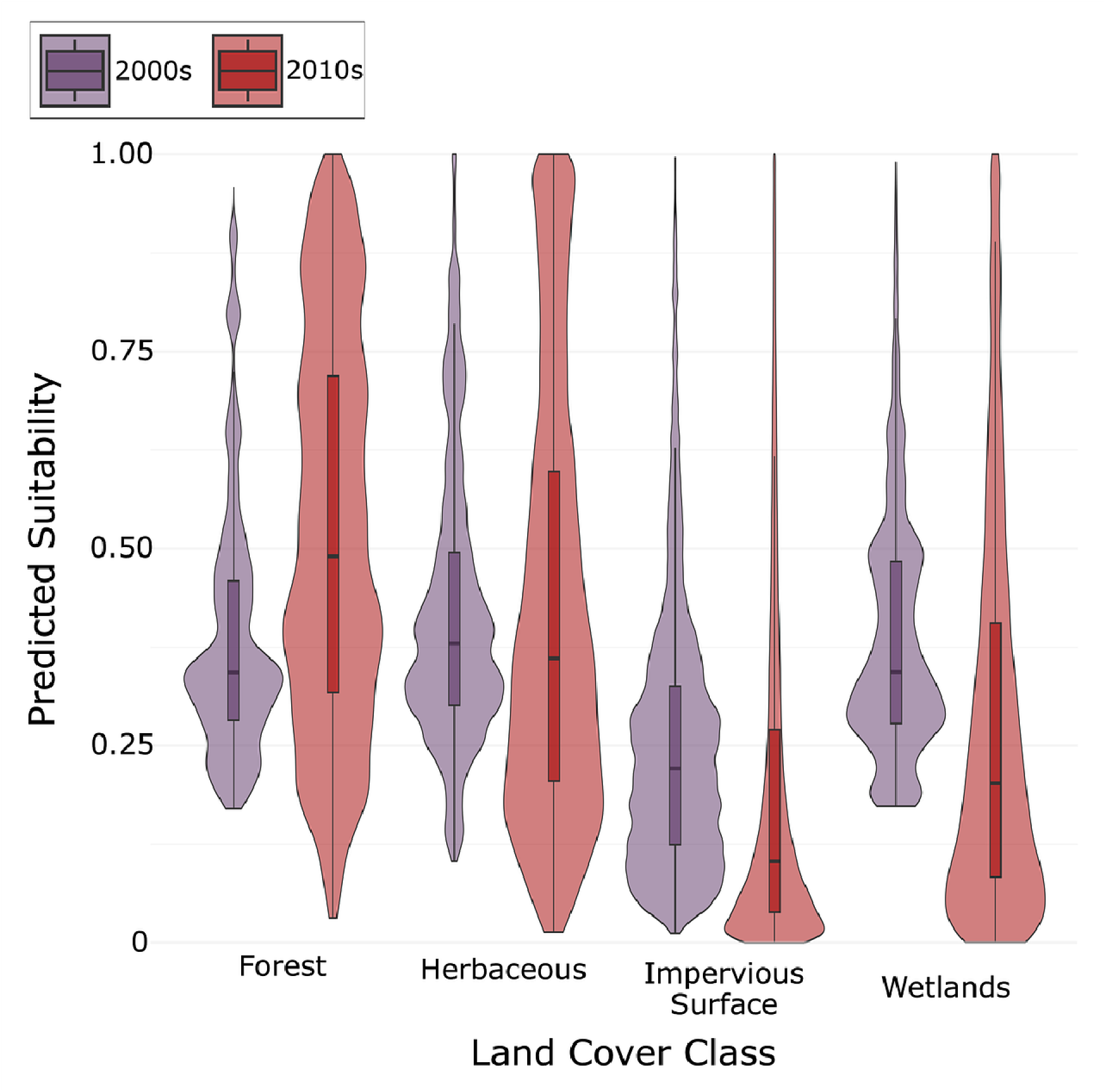
Violin plots of habitat suitability values grouped by land cover class for the 2000s (purple) and 2010s (red). The width of each violin reflects the density of predicted habitat suitability values across the 100 Random Forest SDM iterations.

Herbaceous cover showed minimal change (median difference = –0.019) (Figure 5). The remaining two land cover classes (i.e., cultivated land, open water) were excluded from comparison due to insufficient representation across both decades for a robust analysis.

Partial dependence plots provided insight into the direction and strength of each variable’s effect on suitability across 100 model runs (Figure 6). Solar radiation had a consistently positive relationship with suitability in both decades. Mean temperature showed a weakly positive effect in the 2000s that shifted slightly negative in the 2010s. Maximum temperature followed a similar pattern, with a decline in slope across decades. Minimum temperature was negatively associated with suitability throughout both periods. Vapor pressure deficit had a weak positive relationship in the 2000s but shifted to a negative association in the 2010s.

**Figure 6.**
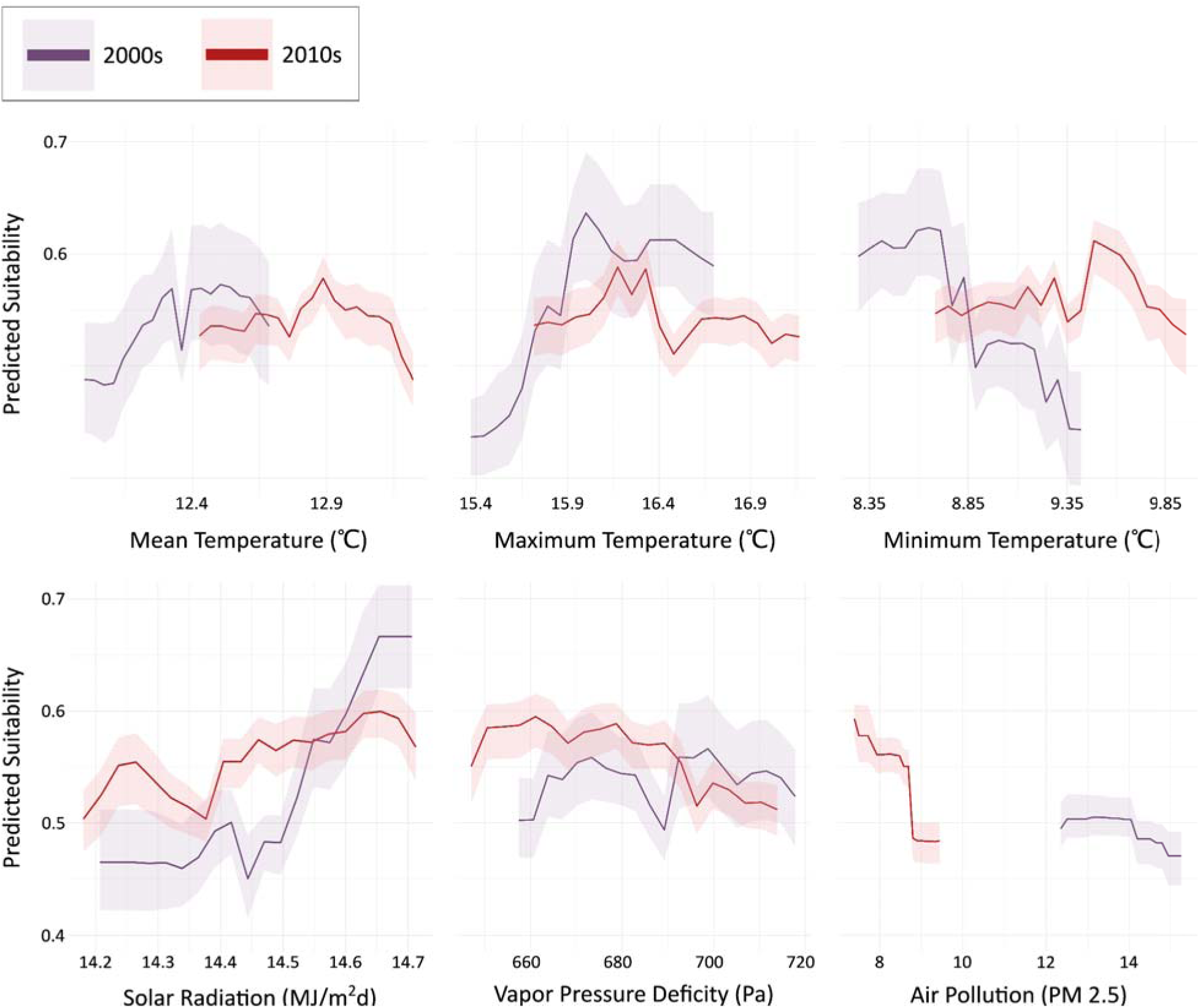
Partial dependence plots for each of the six climatic variables included in Random Forest models. Darker lines indicate the mean across 100 model iterations per decade, while the lighter shaded areas show the full range of predicted values.

## Discussion

Given the widespread and ongoing declines in insect populations worldwide, identifying the primary factors responsible is of urgent scientific concern. In the context of accelerating urban development, it is especially important to assess how urbanization influences habitat suitability for insect populations. To evaluate how urbanization shapes habitat suitability for insects, we used species distribution models (SDMs), powerful computational tools that have been widely used to model species–environment relationships across diverse taxa and habitats, including urban areas (Planillo et al., 2021; Casanelles-Abella et al., 2021). Despite their broad application, SDMs remain underutilized for studying urban invertebrates, particularly diurnal pollinators such as butterflies, bees, and hoverflies. Existing SDM studies on insects have largely focused on invasive species (Koch, 2021, Thomas et al., 2017, Sung et al., 20218), leaving a critical gap in our understanding of how urban environments influence pollinator habitat suitability. Our results revealed a city-wide decline in pollinator habitat suitability in New York City from the 2000s to the 2010s, with pronounced spatial variation across boroughs.

Although we found a net decrease in habitat suitability across New York City’s five boroughs between the 2000s and 2010s models, we also identified localized increases. Notably, suitable habitat for diurnal pollinators between decades increased in Manhattan and in the Bronx, whereas it decreased in the other three boroughs, with the steepest drop in Queens. Although, we cannot rule out plastic or evolved adaptive changes in habitat use. To better understand the drivers of these borough-level shifts in habitat suitability, we examined changes across different land cover classifications and considered the impact of city-wide greening initiatives. Most urban land cover types showed declines in suitability between the 2000s and 2010s; however, urban forest stood out as a notable exception, with a 29.9% increase in suitability, although with much more variation in the 2010s. This positive trend likely reflects major restoration and greening efforts that began around the turn of the century and expanded in the 2010s. For example, Forever Wild, which aimed to protect ecologically valuable land, began in 2001 and was heavily augmented in 2018, adding another 2,500 acres to the program’s extent (Cullman et al., 2023).

Another major program, MillionTreesNYC (2007–2015), aimed to plant one million trees across all five boroughs and met this goal in 2015 (Jones, 2021). These urban forest remediation projects may help explain the increased suitability observed in Manhattan and the Bronx, where many of these initiatives were concentrated. This may also reflect biased urban greening efforts towards areas with higher socioeconomic neighborhoods (Schwarz et al., 2015). Additionally, the Bronx and Manhattan have disproportionately larger amounts of tree canopy for their respective land areas compared to Brooklyn and Queens (Treglia et al., 2021).

Other urban initiatives in New York City may have improved pollinator habitat suitability without directly increasing the urban forest. The Green Infrastructure Program, launched by the NYC Department of Environmental Protection in 2010, expanded throughout the decade and implemented stormwater management features such as green roofs, rain gardens, permeable pavements, and bioswales (Brears, 2023). PlaNYC, a broader sustainability and resilience initiative, further emphasized urban greening, forest remediation, and biodiversity enhancement while setting goals for greenhouse gas reduction and climate adaptation. In addition to ground-level efforts, elevated green spaces became increasingly prominent during this period. The expansion of the High Line Park (Manhattan) throughout the 2010s contributed to habitat creation above street level, and related pedestrian-friendly redesigns, such as tree-lined plazas, bioswales, and expanded sidewalks, prioritized green infrastructure across the city (Aslanoğlu et al., 2025). Finally, financial incentives such as the Green Roof Tax Abatement encouraged private property owners to install green roofs, leading to large-scale installations on public buildings like the Javits Center and in private developments. Together, these initiatives may have played a key role in buffering or reversing declines in pollinator habitat suitability in select boroughs, underscoring the ecological value of sustained urban greening.

While urban greening initiatives may have helped offset declines in some areas, our models suggest that temperature (minimum, mean, and maximum), solar radiation, and vapor pressure are the most influential predictors of habitat suitability. This finding suggests that the dominant environmental pressures on urban pollinators are climate-driven. Temperature and moisture availability are especially critical for insect survival, directly influencing metabolic rates, water loss, and the availability of floral resources, all of which may influence distribution (Neven, 2000; Chown et al., 2011; John et al., 2024; Abdullah, 1961). Rising temperatures and increasing vapor pressure deficits likely contribute to declining habitat suitability by intensifying dehydration stress in insects, reducing pollinator performance (Johnson et al., 2023), and limiting nectar and pollen production in urban vegetation (Scaven & Rafferty, 2013). Greening interventions that increase vegetation cover, reduce impervious surface heat gain, or create shaded microhabitats may help buffer these stressors by moderating temperatures and enhancing local moisture retention (Muñoz & Duarte, 2025; Li et al., 2025; Dimoudi & Nikolopoulou, 2003). Surface solar radiation, which governs light availability via cloudiness, also plays a key role in determining bloom times, microclimate conditions, and pollinator foraging behavior (Watson et al., 2022). These findings underscore that climate remains the dominant limiting factor for urban pollinators.

The variable importance from the order-based models demonstrated highly similar environmental sensitivities between orders, suggesting habitat suitability for all focal orders is determined by similar climatic conditions. This consistency indicates that butterflies, bees, wasps, and beetles respond in comparable ways to key environmental variables, particularly temperature, solar radiation, and vapor pressure. Furthermore, the order-based models closely mirrored the decadal model’s overall trends, reinforcing that no single focal order is disproportionately influencing the broader patterns of change. Given this overlap and the stronger performance of the aggregated model, we rely on the aggregated results as the most comprehensive and robust representation of urban pollinator suitability. Both the 2000s and 2010s aggregated models performed well, with the 2010s model achieving slightly higher evaluation metrics. The aggregated model also outperformed the order-based models overall, likely reflecting increased data availability and improved modeling accuracy in recent years.

While this research offers valuable insights into urban pollinator habitat dynamics, several limitations should be acknowledged. First, our models rely on presence-only data, which can introduce spatial and temporal biases due to uneven sampling effort (Elith & Leathwick, 2009). To mitigate these biases, we implemented multiple corrective strategies, including decade-specific spatial thinning, coordinate uncertainty filtering, and the use of kernel density estimation for background sampling (Steen et al., 2020; Zhang et al., 2018). Second, although our models captured broad environmental trends, they did not account for fine-scale heterogeneity typical of urban landscapes, such as microclimatic variation, floral diversity, soil moisture, and shading from buildings (Cadenasso et al., 2007; Dimoudi & Nikolopoulou, 2003). Incorporating higher-resolution environmental data in future work could improve predictions, especially when paired with species-specific analyses and investigations into how urbanization interacts with other ecological stressors, such as pollution or invasive species. Finally, species distribution models operate under key assumptions, including environmental equilibrium and niche conservatism (Richmond et al., 2010). These assumptions are often challenged in urban settings, where rapid landscape change and human-mediated factors, such as artificial nesting sites, native plant gardens, and beekeeping, can allow species to persist in otherwise unsuitable environments (Fortel et al., 2016; Fukase & Simons, 2016). However, because our models were restricted to New York City, which has been extensively urbanized since the late 19th century, we assume a relative environmental equilibrium within this spatial extent. To further address potential violations of niche conservatism, we employed an order-based modeling approach to capture broader temporal shifts in habitat suitability and reduce taxon-specific biases.

Our models offer valuable insights for both urban pollinator conservation and future city planning. By identifying the most influential environmental variables shaping insect distributions, our models provide a foundation for targeted strategies to support pollinator populations. Efforts to mitigate rising temperatures and improve moisture retention through increased vegetation cover, water-conscious urban design, and climate-adaptive infrastructure could significantly enhance habitat quality for urban insects. These findings can inform the design of green roofs, parks, and other green spaces to better support key invertebrate species, particularly diurnal pollinators that depend on temperature-regulated floral resources. Beyond ecological applications, our urban SDMs also present opportunities for public engagement and outreach. Communicating model results, such as identifying pollinator hotspots or visualizing shifts in habitat suitability, may foster greater public appreciation of urban biodiversity and inspire community-led conservation initiatives. Ultimately, by improving our understanding of how urbanization affects invertebrate biodiversity, this research not only contributes to ecological knowledge, but also supports more informed, resilient, and sustainable urban planning.

## Acknowledgements

We would like to thank New York University for providing the funding needed to make this research possible. We would also like to extend thanks to our colleagues in NYU’s Department of Environmental Studies for their feedback on earlier iterations of the manuscript.

## Declaration of Interest statement

The authors declare no competing financial interests or personal relationships that could have influenced the research presented in this paper.

## Funding

This work was supported by the College of Arts and Sciences, New York University, Dean’s Undergraduate Research Fund.

## Notes

### Competing Interest Statement

The authors have declared no competing interest.

